# EXPA delivery links cell polarity to polarized growth in root hair

**DOI:** 10.64898/2026.11.20.768177

**Authors:** Ke Zhou, Yutong Zheng, Daniel Cosgrove

**Affiliations:** Department of Biology, The Pennsylvania State University, University Park, PA 16802 USA

## Abstract

Turgor pressure, which drives irreversible cell wall enlargement and thereby cell growth, is isotropic. Polarized plant cell growth therefore requires spatially regulated wall-loosening to confine growth to specific domains. These domains are often marked by localized ROP GTPase activity, but how this polarity is translated into localized wall loosening remains unclear. Here, we investigated the delivery of EXPA7, a trichoblast-specific α-expansin that facilitates wall loosening and enlargement during root hair growth. We show that ROP activation directs EXPA7 recruitment to the root hair initiation domain through brefeldin-A-sensitive vesicular trafficking, with YIP4a/b required for its normal secretion and wall-deposition. The recruited EXPA7 contributes to ROP2-induced wall enlargement, and its accumulation accompanies local cell wall bulging. Our findings provide a mechanistic link between ROP activation and cell growth, suggesting how signals converging on ROP activation may control growth.

## Introduction

Spatially regulated wall enlargement underlies polarized plant cell growth and morphogenesis^1^, as exemplified by root hairs^2,3^, cotyledon pavement cells^4,5^, and pollen tubes^6,7^. In these cells, the localized accumulation, membrane anchoring, and activation of plant-specific Rho-of-Plant (ROP) GTPases at the inner leaflet of the plasma membrane establish polar domains that specify the sites of subsequent polarized growth^8,9^. This localized ROP activation is regulated by ROP guanine nucleotide exchange factors (GEFs), which promote GDP release and GTP loading and can themselves be activated through phosphorylation by upstream transmembrane^10^ or cytoplasmic^11^ kinases. Active ROP GTPases, in turn, regulate a broad range of local responses, including local cytoskeletal reorganization^2^, vesicle trafficking, membrane tethering and fusion^12-15^, NADPH oxidase-dependent reactive oxygen species (ROS) production and Ca^2+^ influx^16,17^. At the mechanical level, local cell growth involves sustained, irreversible wall extension (creep) under turgor-generated tensile stress^18,19^. Cellulose microfibrils form the principal tension-bearing network of primary cell walls where noncovalent interactions between microfibrils limit turgor-driven wall creep^20,21^ and interstitial pectin and xyloglucan modulate wall thickness and limit direct microfibril contacts^1,20-23^. How localized ROP activation regulates mechanical processes underlying creep is unclear.

Root hairs provide an excellent model for investigating polarized cell growth. In Arabidopsis, root hair-specific ROP-GEFs accumulate at the root hair initiation domain (RHID) of trichoblasts through upstream mechanisms that remain poorly understood and subsequently recruit and activate ROPs before visible bulge formation^24^. Root hairs then emerge as localized cell wall bulges and elongate by tip growth into tubular extensions^25^. Several cell wall remodeling processes accompany root hair growth. Localized apoplastic acidification has been proposed to induce local acid growth at the RHID^26^. Yet lowering wall pH alone neither triggers root hair initiation nor induces ectopic hair bulging^26^, and how apoplastic acidification leads to local wall expansion is debated^27^. Other cell wall remodeling processes have also been implicated in root hair growth. Localized xyloglucan endotransglycosylase activity accompanies root hair initiation^28^ and RALF22-LRX1/2 complexes organize and condense demethylesterified pectin into periodic circumferential rings during root hair tip growth^29^. How these remodeling processes affect cell wall tensile mechanics and whether they induce sustained wall creep are unclear. Our recent work showed that α-expansin (EXPA) activity is required for root hair tip growth^30^. EXPAs promote wall creep by disrupting noncovalent interactions between cellulose microfibrils^18,31^, providing a potential mechanical explanation for root hair growth.

Canonical α-expansins from distinct phylogenetic clades, including the trichoblast-specific EXPA7, can rescue the root-hairless phenotype caused by EXPA deficiency and specifically accumulate in the cell walls of root hair bulges^30^. This spatial coincidence suggests that localized EXPA deposition may promote root hair outgrowth by inducing wall creep. In this study, we examined how ROP2 activation regulates EXPA7 trafficking and secretion. We also used pharmacological and genetic perturbations of vesicle trafficking to dissect its polarized delivery pathway.

## Results

### EXPA7 is polarly delivered to the root hair initiation domain

We previously showed that trichoblast-specific EXPA7 exhibits characteristic cell wall association and polar enrichment at the apices of root hair bulges and elongating root hairs^30^. Here, we used time-lapse imaging to track EXPA7-mCherry localization throughout root hair development (**Mov. 1**). Fluorescence can be first detected at the RHID before a distinct bulge formed (**Fig. 1A**) and progressively intensified as the bulge developed (**Fig. 1B-C**). These observations indicate that EXPA7 accumulation at the RHID coincides with the onset of local cell wall extension that gives rise to root hair bulging and persists throughout root hair initiation.

**Fig. 1.**
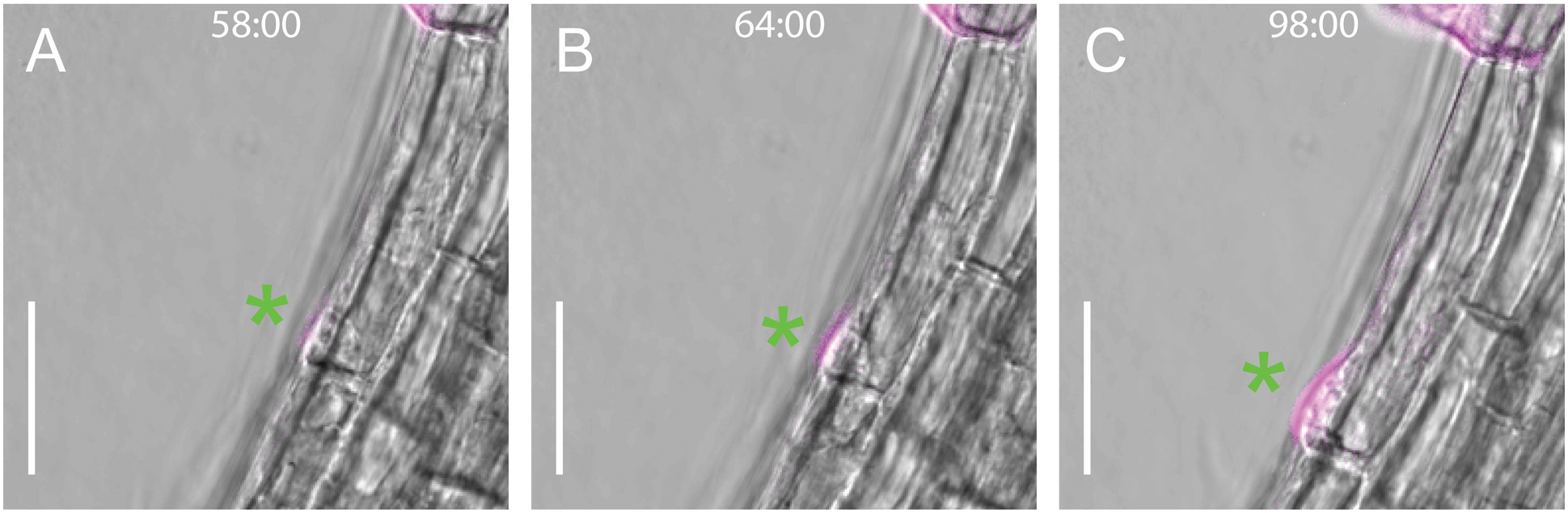
EXPA7 accumulation accompanies hair bulging. (**A-C**) Time-lapse confocal images from Mov.1 showing a trichoblast expressing *EXPA7pro::EXPA7-mCherry*. EXPA7-mCherry fluorescence (magenta) is overlaid on bright-field images. Green asterisks mark the same root hair initiation domain (RHID) indicated by the green asterisk in Mov.1. Fluorescence at the RHID is first detectable at 58 min (A), before a distinct bulge forms, and progressively intensifies as the bulge develops at 64 min (B) and 98 min (C). Times indicate elapsed time from the start of Movie 1. Scale bars, 20 μm.

In Arabidopsis, the polarity that positions root hair at the rootward end of differentiated trichoblast depends on the auxin gradient along their shootward-rootward axis^25,32^. Low-dose brefeldin A (BFA; 10 µM) inhibits hair elongation and shifts the positioning of newly formed root hairs, likely by interfering with auxin influx^32^. We transferred seedlings expressing EXPA7-mCherry under the trichoblast-specific *EXPA7* promoter to ½MS medium containing 10 µM BFA for overnight treatment. EXPA7 accumulated at newly formed root hair bulges (**Fig. 2A-2B**), despite BFA-induced shifts in their positioning (**Fig. 2B-2C**). This suggests that the polarity mechanism specifying root hair positioning recruits EXPA7.

**Fig. 2.**
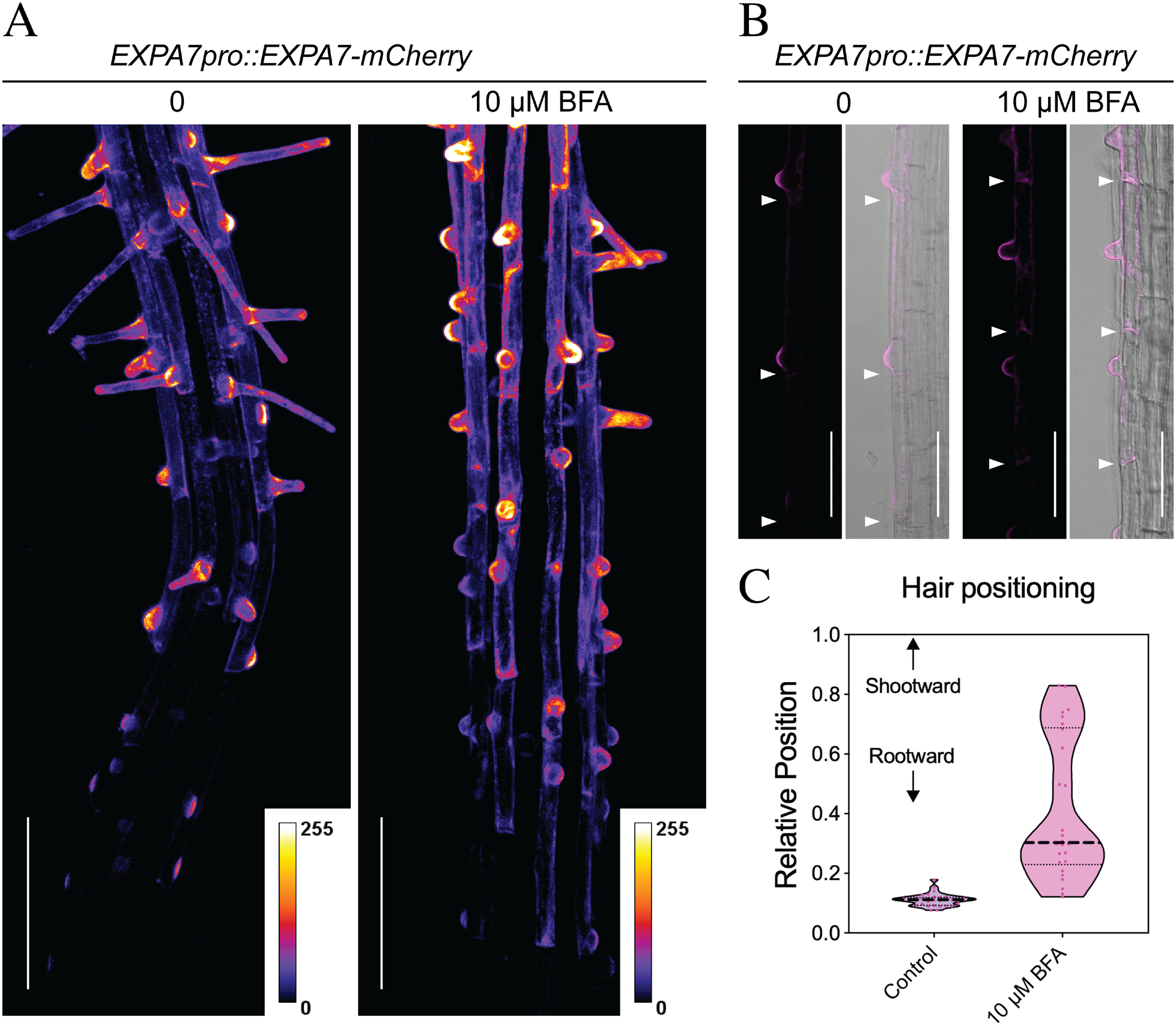
Polarity specifying hair-positioning recruits EXPA7. Four-day-old seedlings expressing *EXPA7pro::EXPA7-mCherry* were transferred to ½MS medium with or without 10 μM brefeldin A (BFA) and incubated overnight. (**A**) Maximum-intensity projections of confocal z-stacks showing EXPA7-mCherry distribution in early differentiation zones of control (left) and BFA-treated (right) roots. Fluorescence is displayed using the Fire pseudocolor lookup table; color scales indicate fluorescence intensity (0-255). (**B**) Representative images of EXPA7-mCherry fluorescence (magenta) and corresponding bright-field overlays in control (left) and BFA-treated (right) roots, showing BFA-induced shifts in the positioning of newly formed root hair bulges. White arrowheads mark cell boundaries. (**C**) Relative positions of newly formed root hairs within trichoblasts. Relative position was calculated as the longitudinal distance from the rootward cell boundary to the center of the root hair base, normalized to trichoblast length. Values of 0 and 1 indicate the rootward and shootward cell boundaries, respectively. Each point represents a root hair; n = 31 (control) and 26 (BFA). Scale bars, 100 µm (A) and 50 µm (B).

We next expressed EXPA7-mCherry before root hair initiation using the *PIN2* promoter in elongation-zone epidermal cells^33^. *PIN2pro::EXPA7-mCherry* was introduced into the *PIN2pro::PIN2-GFP* line, visualizing their distinct distributions. EXPA7-mCherry exhibited negligible polar enrichment, whereas PIN2-GFP retained its characteristic polar distribution at the shootward (apical) plasma membrane of root epidermal cells (**Fig. S1A-S1C**). Transmembrane PIN2, an auxin efflux carrier, maintains its polar distribution through polarized secretion, spatially regulated endocytosis, and recycling^34^. The contrasting distributions of PIN2 and EXPA7 within the same cells suggest that their polar localization relies on distinct targeting mechanisms and that EXPA7 recruitment to the RHID requires machinery specifically associated with trichoblast differentiation. Consistently, this early EXPA7 expression without polar enrichment neither advanced root hair initiation (**Fig. S1D**) nor shifted hair positioning (**Fig. S1E**).

Together, these results suggest that EXPA7 recruitment to the RHID is directed by the pre-established polarity specifying root hair position in trichoblasts and accompanies root hair bulging.

### EXPA7 delivery requires active ROP2

Distinct ROP-GEFs act sequentially to recruit and activate ROP at the RHID; loss of early-expressed *ROP*-*GEF3* and *ROP*-*GEF4* delayed hair initiation^24^ (**Fig. 3A-3C**). At the early differentiation zone (brackets in **Fig. 3A-3B**), the RHID-accumulation of EXPA7-mCherry observed in *Col-0* (**Fig. 3D**) was absent in the *gef3/gef4* mutant (**Fig. 3E**), despite comparable EXPA7-mCherry abundance between genotypes, as assessed by immunoblotting (**Fig. S2A-2B**). Plasmolyzed trichoblast showed pronounced intracellular accumulation of EXPA7-mCherry in *gef3/gef4* (**Fig. 3F-3I**), suggesting that impaired ROP activation strongly compromised EXPA7 delivery to RHID.

**Fig. 3.**
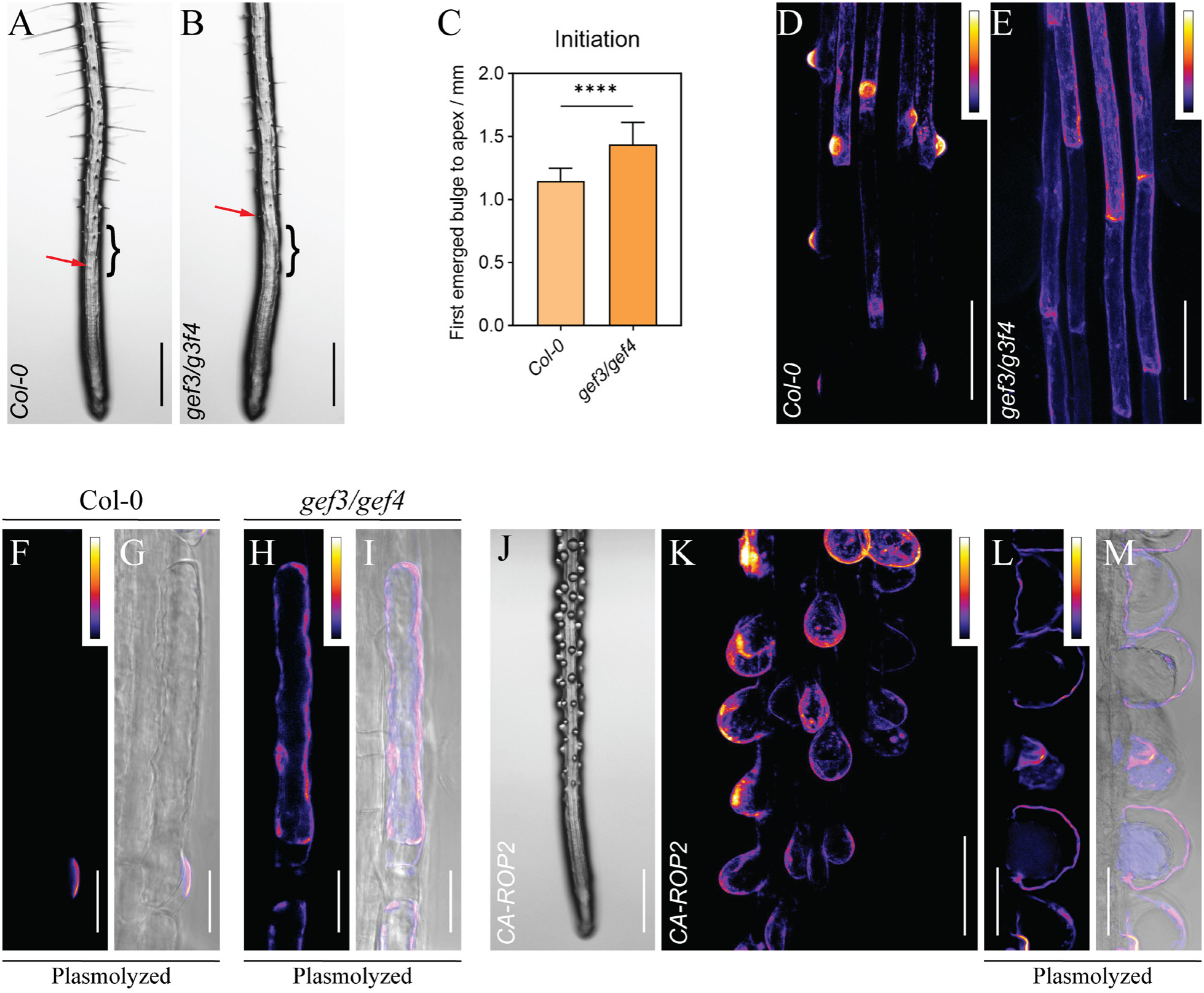
ROP activation regulates EXPA7 delivery. *EXPA7pro::EXPA7-mCherry* construct was introduced into Col-0, *gef3/gef4* and *EXPA7pro::CA-ROP2* lines. (**A** and **B**) Representative root images of Col-0 (A) and *gef3/gef4* (B) seedlings. Red arrows indicate the first visible root hair bulge; brackets indicate the early differentiation zone. (**C**) Distance from the root apex to the first visible root hair bulge in Col-0 and *gef3/gef4* (n=12 and 14 seedlings, respectively). Data are means ± s.d; ****, P<0.0001 (t-test). (**D** and **E**) Maximum-intensity projections of confocal z-stacks showing EXPA7-mCherry distribution in the early differentiation zone of Col-0 (D) and *gef3/gef4* (E). (**F**-**I**) EXPA7-mCherry localization following plasmolysis in Col-0 (F, G) and *gef3/gef4* (H, I). Fluorescence images (F, H) and overlays with bright-field images (G, I) show wall-associated fluorescence in Col-0 and intracellular accumulation in *gef3/gef4*. (**J**) Representative root image showing spherical root hairs induced by *EXPA7pro::CA-ROP2*. (**K**) Maximum-intensity projection of a confocal z-stack showing EXPA7-mCherry distribution in bulbous root hairs expressing CA-ROP2. (**L** and **M**) EXPA7-mCherry localization in CA-ROP2-expressing root hairs following plasmolysis. The fluorescence image (L) and overlay with the bright-field image (M) show fluorescence broadly distributed along the cell walls. Fluorescence images are displayed using the Fire pseudocolor lookup table; color bars indicate relative fluorescence intensity (0-255). Scale bars, 500 μm (A, B, and J), 100 μm (D, E, and K), 20 μm (F-I), and 50 μm (L and M).

Conversely, a Gly-to-Val substitution maintains ROP2 in a constitutively active, membrane-associated state (CA-ROP2), broadening the polar domains and induces swelling at sites of polarized growth^2,3^. Expression of CA-ROP2 under the trichoblast-specific *EXPA7* promoter produced spherical root hairs lacking a distinct apex (**Fig. 3J)**. EXPA7-mCherry showed widespread accumulation in these bulbous hairs (**Fig. 3K**), in marked contrast to its localized accumulation at bulges and growing tips in wild-type root hairs^30^. Following plasmolysis, EXPA7-mCherry was shown broadly distributed along their spherical cell walls (**Fig. 3L-3M**).

Impaired ROP activation abolishes EXPA7 accumulation at the RHID, whereas constitutive ROP2 activation broadens its distribution, linking the spatial pattern of EXPA7 accumulation to ROP activation.

### EXPA7 contributes to ROP2-induced localized wall enlargement

We then introduced *EXPA7pro::CA-ROP2* into *expa7/expa18* lines expressing either *EXPA7pro::EXPA7-mCherry* or *EXPA7pro::EXPA7D136N-mCherry*; the latter encodes the active-site D136N mutant previously shown to lack wall-creep activity and fail to rescue root hair growth in *expa7/expa18*^30^ (**Fig. 4A-4B**). Despite comparable CA-ROP2 expression levels (**Fig. S3**), hairs in the *EXPA7-mCherry* line had markedly larger diameters (**Fig. 4E**) and correspondingly greater surface areas (**Fig. 4F**). These results indicate that EXPA7-induced wall creep contributes to wall enlargement driven by ROP2 activation.

**Fig. 4.**
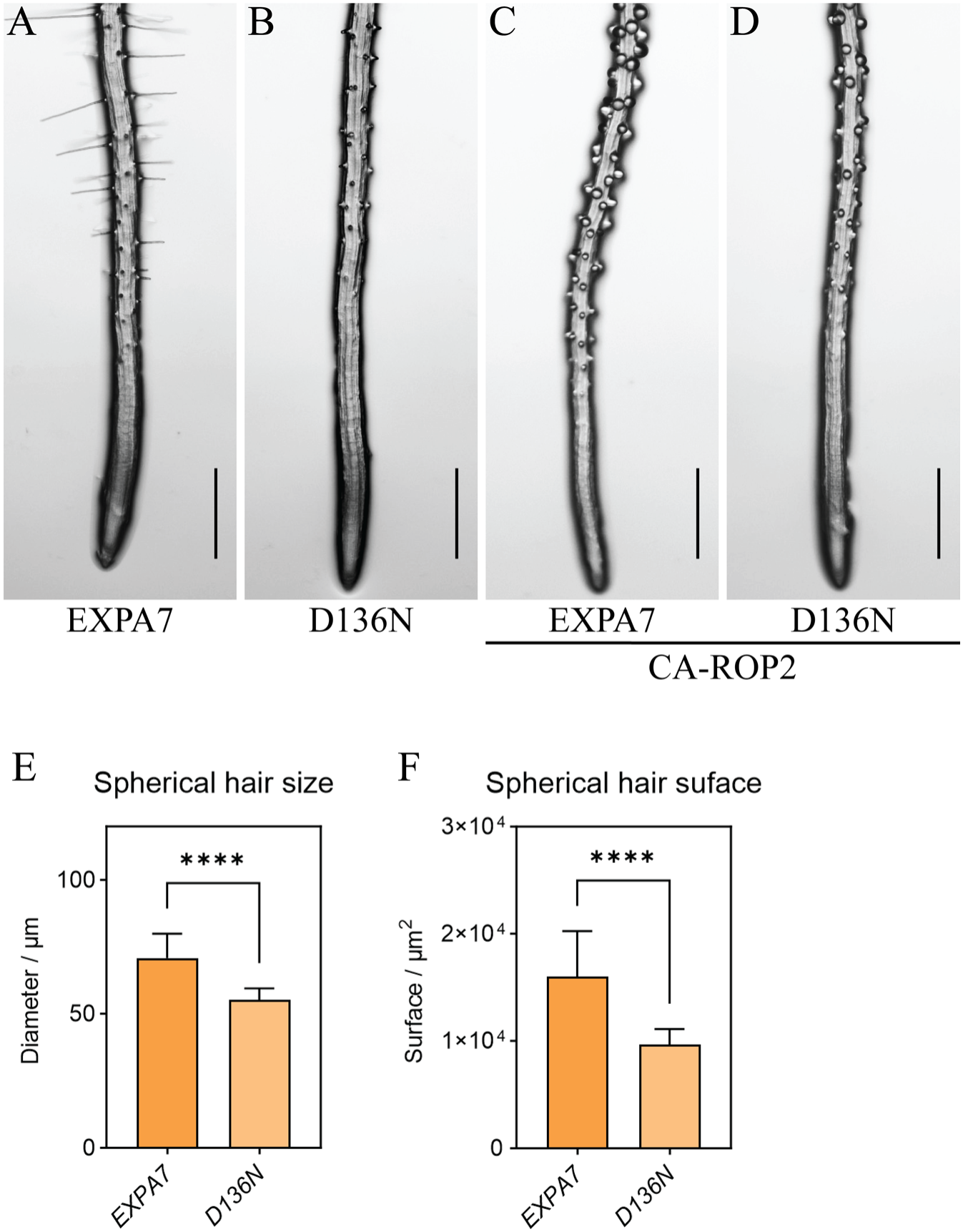
EXPA7 contributes to CA-ROP2-induced wall enlargement. *EXPA7pro::CA-ROP2* was introduced into *expa7/expa18* lines expressing either functional EXPA7-mCherry or the inactive EXPA7D136N-mCherry variant under the *EXPA7* promoter. (**A-D**) Representative root images of the EXPA7-mCherry (A), EXPA7D136N-mCherry (B), CA-ROP2/EXPA7-mCherry (C), and CA-ROP2/EXPA7D136N-mCherry (D) lines. (**E** and **F**) Diameter (E) and estimated surface area (F) of spherical root hairs in the EXPA7-mCherry and EXPA7D136N-mCherry lines expressing CA-ROP2. Measurements were averaged over the 10 largest spherical root hairs per root; n = 12 roots for each. Data are means ± s.d; ****, P<0.0001 (t-test). Scale bars, 500 μm (A-D).

### EXPA7 transport requires brefeldin-A-sensitive vesicular trafficking

ROP activation regulates vesicle trafficking, membrane tethering and fusion^12-15^. We therefore investigated the trafficking pathway underlying ROP-dependent EXPA7 delivery. At the trans-Golgi network (TGN), ARF-GEF-mediated activation of ARF GTPases promotes vesicle formation and trafficking^35,36^. High-dose BFA (50 µM) binds and inhibits BFA-sensitive ARF-GEFs^37^, blocking vesicle trafficking and aggregating secretory cargo within intracellular membrane structures termed BFA bodies^38^. Seedlings expressing *EXPA7pro::EXPA7-mCherry* were incubated in liquid ½ MS medium containing 50 µM BFA for 1 hour and we observed fluorescence accumulation in cytoplasmic BFA bodies (**Fig. 5A-5D**), supporting the involvement of BFA-sensitive vesicular secretion in EXPA7 delivery.

**Fig. 5.**
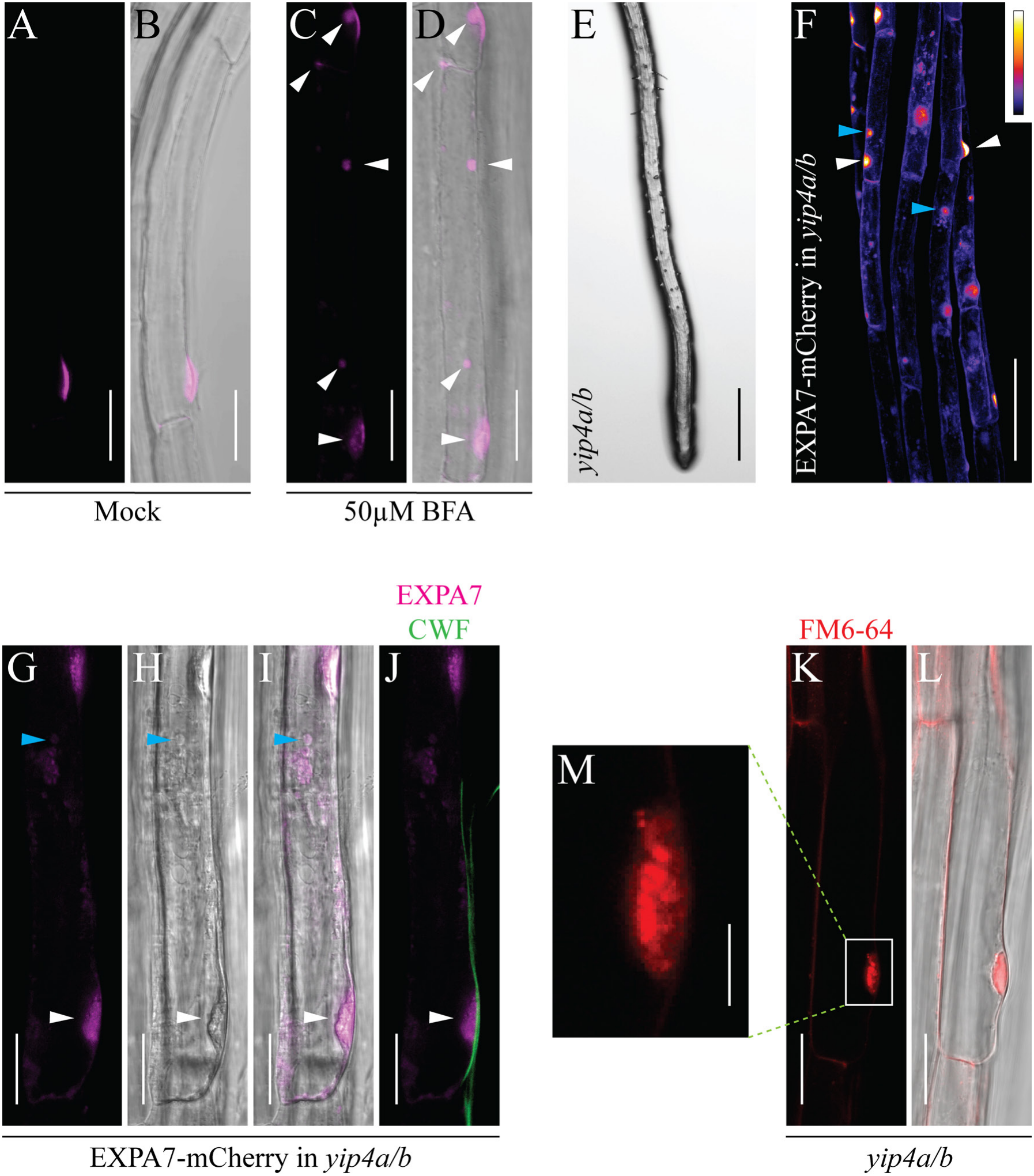
YIP4a/b contribute to EXPA7 secretion into apoplast. (**A-D**) Confocal images of EXPA7-mCherry (*EXPA7pro::EXPA7-mCherry*) in mock-treated seedlings (A and B) and seedlings treated with 50 µM BFA for 60 min (C and D). EXPA7-mCherry fluorescence (magenta; A and C) and corresponding bright-field overlays (B and D) are shown. White arrowheads indicate BFA bodies (C and D). (**E**) Representative root image of a *yip4a/b* seedling root tip, showing impaired root hair elongation. (**F**) Confocal image showing intracellular accumulation and retained enrichment of EXPA7-mCherry at the RHID in *yip4a/b*. Green and white arrowheads mark EXPA7-mCherry accumulation outside and at the RHID, respectively. Fluorescence is displayed using the Fire pseudocolor lookup table; the color bar indicates relative fluorescence intensity. (**G-J**) EXPA7-mCherry fluorescence (magenta; G), the corresponding bright-field image (H) and overlay (I), and a merged image of EXPA7-mCherry (magenta) and calcofluor white (CFW; green) fluorescence (J), showing EXPA7-mCherry accumulation in an aberrant shaped structure beneath the cell wall at the RHID in *yip4a/b*. Cyan arrowheads indicate intracellular EXPA7-mCherry accumulation, and white arrowheads indicate the aberrant shaped structure. (**K**-**L**) FM4-64 fluorescence (red; K) and the corresponding bright-field overlay (L), showing a similar aberrant shaped structure in control *yip4a/b* trichoblast. (**M**) Enlarged view of the boxed region in K. Scale bars, 500 µm (E), 100 µm (F), and 20 µm (A-D and G-L), and 5 µm (M).

### EXPA7 secretion into the apoplast requires YIP4a/b

ROP-driven root hair growth requires the TGN-localized YPT/RAB GTPase-interacting proteins 4a and 4b (YIP4a/b), which also contribute to active ROP accumulation at the RHID^39,40^. In the *yip4a/b* mutant, root hair initiation and elongation are severely impaired^41^ (**Fig. 5E**). We therefore introduced *EXPA7pro::EXPA7-mCherry* into the *yip4a/b* mutant. Notably, in contrast to the loss of accumulation at the RHID in *gef3/gef4* (**Fig. 3H-3I**), EXPA7-mCherry in *yip4a/b* showed both intracellular accumulation and retained enrichment at the RHID (**Fig. 5F**), suggesting that RHID polarity is established but attenuated, consistent with the persistence of smaller, less intense, active ROP2 patches in this mutant^41^.

However, the retained accumulation of EXPA7 at the RHID (**Fig. 5F**) appears insufficient to support root hair elongation in *yip4a/b* (**Fig. 5E**). Closer observation on trichoblast at the early differentiation zone revealed EXPA7-mCherry accumulation within aberrant structures at the RHID beneath the cell wall stained by calcofluor white (**Fig. 5G-5J**). Similar structures were also observed in control *yip4a/b* trichoblast and were stained with the membrane dye FM4-64 (**Fig. 5K-5M**), suggesting that they might represent aggregates of vesicles that failed to fuse with the target membrane and release their cargo into the apoplast.

YIP4a/b are required for proper EXPA7 release into the apoplast but are not strictly required for its polar enrichment at the RHID, suggesting that they may contribute to EXPA7-contained vesicle fusion with the target membrane.

### ROP-dependent EXPA recruitment may vary across cell types

In addition to root hair growth, active ROP GTPases regulate other forms of polarized growth through distinct signaling modules: ROP2/4/6 regulate pavement cell morphogenesis^4,42^, whereas ROP1/3/5/9 participate in pollen tube growth^6,7,43^. We expressed EXPA7-mCherry under the constitutive *UBQ10* promoter (*UBQ10pro::EXPA7-mCherry*) and observed EXPA7 distribution in these cell types. EXPA7-mCherry accumulated at pavement cell lobes, in a pattern consistent with the reported distribution of active ROP2^44^ (**Fig. S4A**). In pollen tubes, however, it did not show apical enrichment, and fluorescence was predominantly cytoplasmic (**Fig. S4B**-**S4C**). Given the functional diversity of EXPAs^30^ and ROPs^45^, these contrasting distributions suggest that the mechanism by which ROPs promote polarized growth through EXPA delivery may vary across cell types.

## Discussion

Spatially controlled cell expansion underlies plant cell morphogenesis^1^. Our findings identify polarized EXPA7 delivery as a physical link between ROP2-defined cell polarity and localized growth during root hair development. This mechanism extends the acid-growth framework by adding spatial control of expansin availability to pH-dependent activation. The functional redundancy and diversity within the expansin superfamily^30^ and ROP family^46^ suggest that this mechanism may be broadly conserved yet functionally diverse.

ROP activation can regulate vesicle targeting by reorganizing the actin and microtubule networks^4,15,42^, and engage the machinery for vesicle tethering^14^ and membrane fusion^13^. Our observation that EXPA7 trafficking is BFA-sensitive is consistent with a vesicular route for its delivery. Notably, TGN-localized YIP4a/b help coordinate secretory trafficking and maintain the TGN localization of the SNARE SYP61^39,47^, which physically associates with the plasma membrane SNARE SYP121^48^, a direct binding partner of active ROP2^13^. Whether SYP61 and SYP121 mediate EXPA7 secretion remains to be established. Nevertheless, EXPA7 accumulation in aberrant membrane structures beneath cell wall at the RHID in *yip4a/b* suggests that YIP4a/b may act at a late step in secretion, potentially involving membrane fusion and cargo release. Together, these observations suggest that ROP activation may coordinate multiple steps of vesicular trafficking to promote polarized EXPA7 recruitment.

ROP GTPase act as molecular switches that integrate diverse upstream signals^45^. The 11 structurally conserved Arabidopsis ROPs have distinct functions^46^, including ROP11-mediated secondary wall patterning in xylem vessels^49^, and ROP10-mediated inhibition of abscisic acid (ABA) responses^50^. Several kinase pathways converge on ROP activation. For example, the cell-surface receptor kinase PRK2 may relay extracellular cues during pollen tube growth by phosphorylating and activating ROP-GEF1, which in turn activates ROP1^10^. Similarly, the receptor kinase FER links cell wall integrity cues to ROP2 activation in root hairs and ROP6 activation in pavement cells by binding and potentially phosphorylating ROP-GEFs^17,51^. Auxin may signal through cell-surface transmembrane kinases (TMKs)^52^ to the receptor-like cytoplasmic kinases (RLCKs) RAK1-4, which phosphorylate and activate ROP-GEF1, thereby activating ROP GTPases^11^. In root hairs, positive feedback sustains local ROP activity: ROP effectors recruit AGC1.5 subfamily kinases that phosphorylate ROP-GEF4/10 and maintain their polar targeting^53^. ROP-directed α-expansin delivery may therefore provide a shared downstream route linking diverse kinase signals to localized growth.

## Supporting information

MOV. 1

## Acknowledgments

We are grateful to Professor Klaus Palme (University of Freiburg) for his thoughtful discussions and constructive feedback on an earlier version of this manuscript. We thank Heather McFarlane (University of Toronto) for kindly providing the *yip4a/b* mutant seeds used in this study.

## Author contributions

Conceptualization: KZ

Methodology: KZ

Investigation: KZ, YZ

Visualization: KZ

Funding acquisition: DJC

Project administration:KZ

Supervision: KZ, DJC

Writing – original draft: KZ

Writing – review & editing: DJC

## Data, code, and materials availability

All data are available in the main text or the supplementary materials.

## Materials and Methods

### Plant materials and growth conditions

For root hair observation, seeds were surface-sterilized and sown on half-strength Murashige and Skoog (½ MS) agar plates (pH 5.8; 0.5 g L^−1^ MES adjusted with 1 M KOH) supplemented with 1% (w/v) sucrose and 0.5% (w/v) Gelzan (G-1910, Sigma). After stratification in darkness at 4 °C for 3 days, plates were positioned vertically, and seedlings were grown at 22 °C under white light (100 µmol m^-2^ s^-1^) with a 16h light/8h dark photoperiod for 4 days before imaging. For plasmolysis, roots were treated with 500 mM mannitol.

The *expa7/expa18* mutant and transgenic lines expressing EXPA7-mCherry or EXPA7D136N-mCherry were described recently^30^. The *yip4a/b* mutant was kindly provided by Heather McFarlane^39^. The *gef3-1* (SALK_079879C) and *gef4-2* (SALK_107520C) mutants^24^ were obtained from the Arabidopsis Biological Resource Center (ABRC). A *PIN2pro::PIN2-GFP* reporter line co-expressing *PIN2pro::EXPA7-mCherry* was used to visualize the subcellular distributions of PIN2-GFP and EXPA7-mCherry. A 538-bp fragment upstream of the EXPA 7 coding region was used as the EXPA7 promoter, as described previously^30^. A 1.3-kb fragment upstream of the *PIN2* coding region was used as the *PIN2* promoter^54^. Constitutively active ROP2 (CA-ROP2) was generated by site-directed mutagenesis to replace glycine at position 14 with valine (G14V)^2^. A 634-bp fragment immediately upstream of the *UBQ10* start codon was used as the *UBQ10* promoter^55^.

All constructs involving ROP2 were generated in the pEGAD^56^ binary vector (ABRC), which confers Basta resistance and places GFP at the N terminus of ROP2^2^. All constructs involving EXPA7 were generated using the modified pCAMBIA binary vector described in our previous study, which contains a plant-codon-optimized mCherry sequence for C-terminal tagging and a hygromycin resistance marker^30^. All constructs were introduced into *Agrobacterium tumefaciens* strain GV3101, and Arabidopsis plants were transformed using the floral dip method.

### Imaging

Seedling roots used for root hair analysis were imaged using an Olympus MVX10 Macro Zoom Microscope. For subcellular localization analysis, samples were imaged using a Zeiss LSM780 Axio Observer laser scanning confocal microscope. Excitation wavelengths and emission detection ranges were 561 nm and 580-652 nm for mCherry, 488 nm and 500-550 nm for GFP, 561 nm and 650-750 nm for FM4-64, and 405 nm and 425-475 nm for CFW, respectively. Maximum-intensity projections of confocal z-stacks were generated using ZEISS ZEN 3.8 software.

Where indicated, fluorescence images are displayed using the Fire LUT by ImageJ (1.53u version) to facilitate visualization of relative fluorescence intensity patterns. In this pseudocolor scale, pixel intensities are mapped from low to high signal, with darker colors representing lower relative fluorescence and warmer colors representing higher relative fluorescence. The accompanying color bars indicate the displayed 8-bit intensity range from 0 to 255. These pseudocolor images are more informative for visualization of fluorescence distribution compared with absolute fluorescence quantification.

Quantification of spherical root hair size: for each root, the diameters of the 10 largest spherical root hairs were measured in ImageJ. Surface area was estimated for each root hair by approximating it as a sphere and applying the formula A = 4πr^2^, where r is half the measured diameter. Mean diameter and estimated surface area were calculated for each root and used for statistical analysis across roots, with each root treated as an independent biological replicate.

### Western-blotting

For western blot analysis, four-day-old seedling roots were collected, and ground in PBS buffer (pH7.4) containing protease inhibitors (Sigma, cOmplete, 11836170001). The extracts were mixed with 4× SDS loading buffer containing 5 mM DTT and boiled at 95°C for 5 min. Prior to immunoblotting, protein loading was adjusted based on 10% SDS-PAGE to ensure comparable loading among samples. EXPA7-mCherry and GFP-CA-ROP2 were detected using anti-mCherry (Sigma, SAB4702107) and anti-GFP (invitrogen, MA5-15256) primary antibodies, respectively, followed by a Sheep Anti-Rabbit (Rockland, 611-6502)/Goat Anti-mouse (Sigma, A6163) IgG alkaline phosphatase (AP)-conjugated secondary antibody.

---

**Fig. S1.**
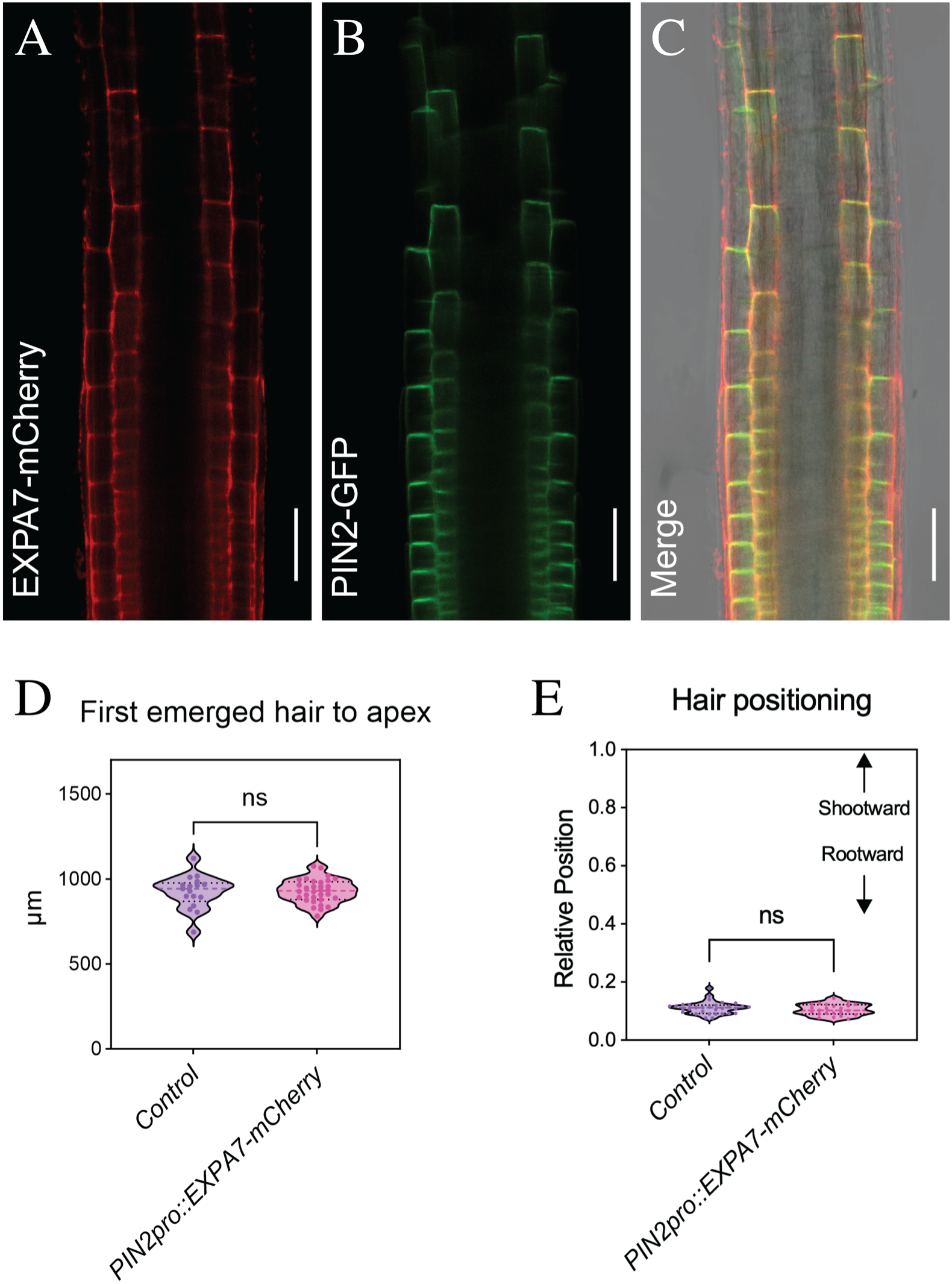
EXPA7-mCherry localization in elongation zone epidermal cells. Four-day-old seedling co-expressing *PIN2pro::EXPA7-mCherry* and *PIN2pro::PIN2-GFP* were observed. (**A-C**) Representative confocal images of epidermal cells in the elongation zone. EXPA7-mCherry fluorescence (red; A), PIN2-GFP fluorescence (green; B), and an overlay of both channels with the corresponding bright-field image (C) are shown. (**D**) Distance from the root apex to the first visible root hair bulge in control and *PIN2pro::EXPA7-mCherry* seedlings (n=17 and 32 seedlings, respectively). (**E**) Relative positions of mature root hairs within trichoblasts, calculated as the longitudinal distance from the rootward cell boundary to the center of the root hair base, normalized to trichoblast length. Values of 0 and 1 indicate the rootward and shootward cell boundaries, respectively (n = 31 and 23 root hairs for control and *PIN2pro::EXPA7-mCherry*, respectively). Each point represents a seedling (D) or a root hair (E). Violin plots show the distributions of individual measurements. Statistical significance was assessed using a t-test; ns, not significant. Scale bars, 50 µm (A-C).

**Fig. S2.**
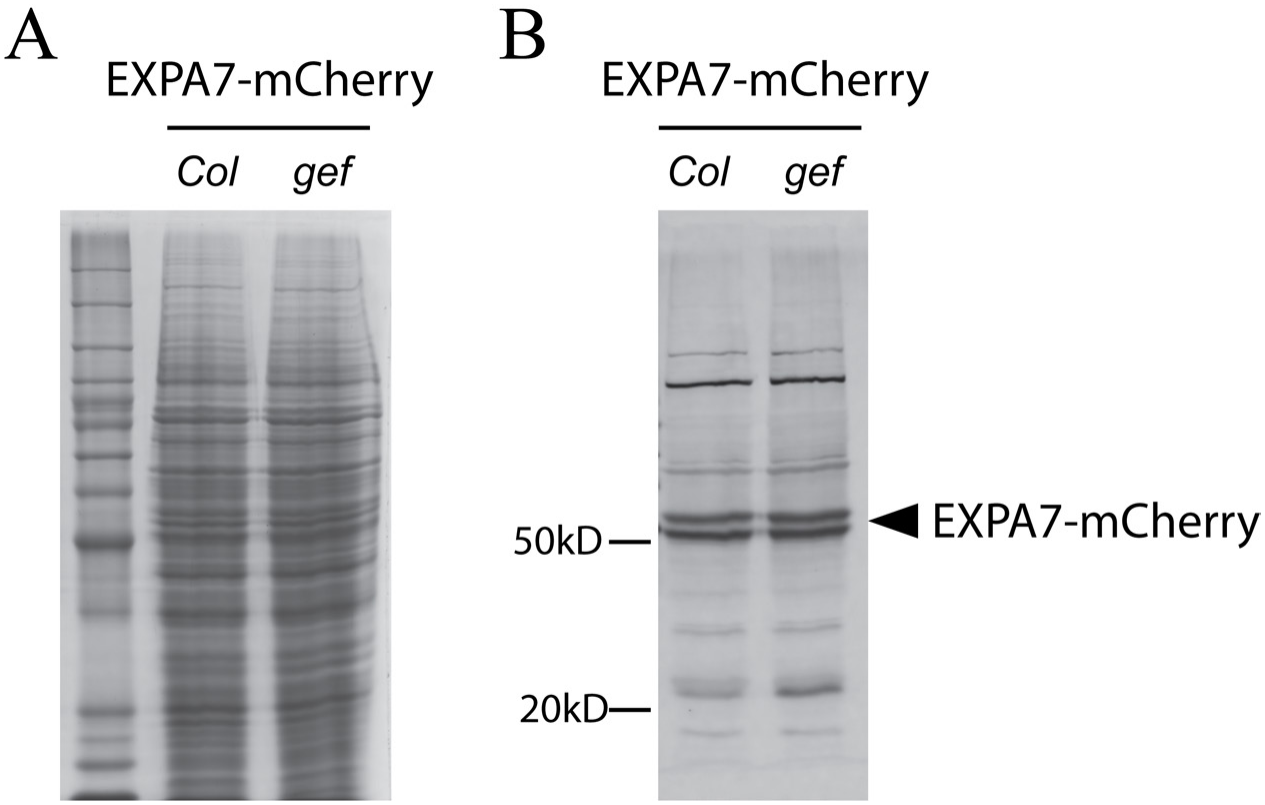
EXPA7-mCherry protein abundance is comparable in Col-0 and *gef3/gef4*. **(A)** Total protein staining of root extracts from 4-day-old Col-0 and *gef3/gef4* seedlings expressing *EXPA7pro::EXPA7-mCherry*, shown as a loading control. **(B)** Immunoblot analysis of EXPA7-mCherry in the corresponding protein extracts using an anti-mCherry antibody. The black arrowhead indicates the EXPA7-mCherry band (predicted molecular mass of the mature fusion protein, approximately 53 kDa), whose identity was established previously^30^.

**Fig. S3.**
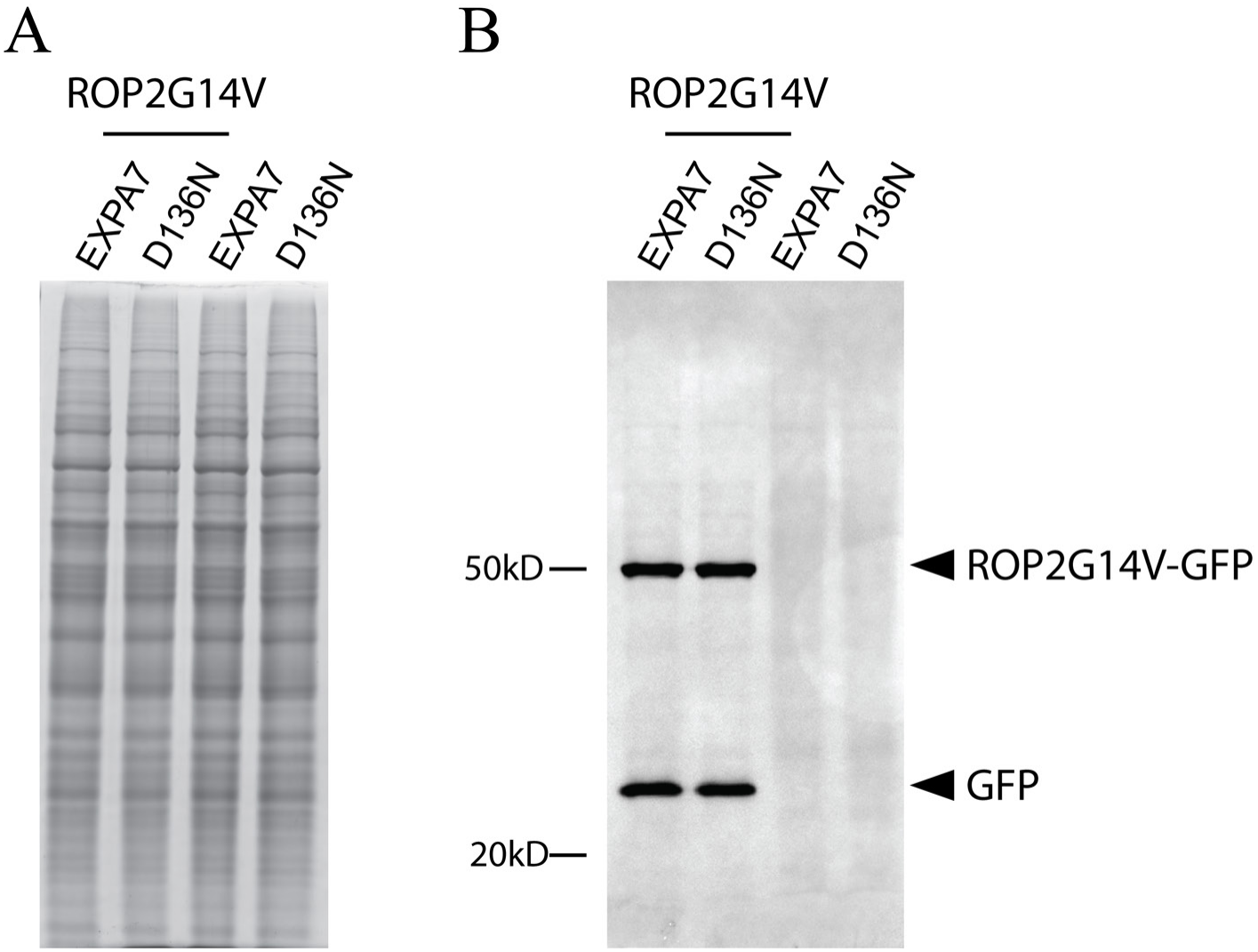
ROP2-G14V protein abundance is comparable between EXPA7-mCherry and EXPA7D136N-mCherry lines. GFP-tagged ROP2-G14V was expressed under the EXPA7 promoter in previously characterized EXPA7-mCherry and EXPA7D136N-mCherry lines, both in the *expa7/expa18* background. (**A**) Total protein staining of root extracts from 4-day-old seedlings of the two lines, shown as a loading control. (**B**) Immunoblot analysis of GFP-tagged ROP2-G14V in the corresponding protein extracts using an anti-GFP antibody. The black arrowhead indicates the GFP-tagged ROP2-G14V fusion protein (predicted molecular mass, approximately 49 kDa).

**Fig. S4.**
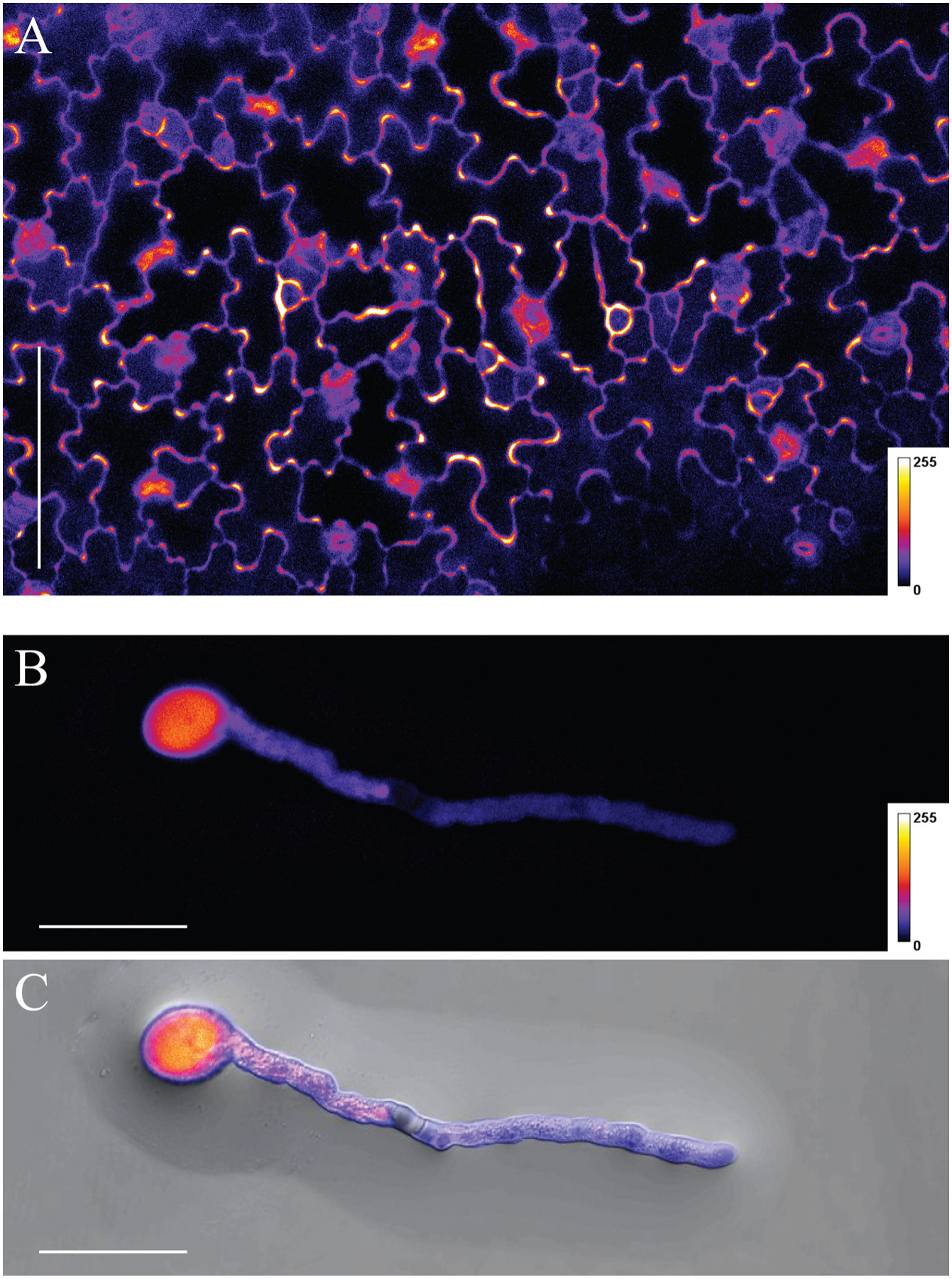
EXPA7-mCherry accumulates at pavement cell lobes but not pollen tube apex. EXPA7-mCherry was expressed under the constitutive UBQ10 promoter (*UBQ10pro::EXPA7-mCherry*). (**A**) Representative confocal image of cotyledon pavement cells from 3-day-old seedlings, showing EXPA7-mCherry accumulation at cell lobes. (**B** and **C**) EXPA7-mCherry distribution in a pollen tube following 10 h of *in vitro* pollen culture. (B) Confocal fluorescence image; (C) overlay with the corresponding bright-field image. Fluorescence is predominantly cytoplasmic and shows no apical enrichment. EXPA7-mCherry fluorescence is displayed using the Fire pseudocolor lookup table in all panels; color bars indicate the displayed fluorescence intensity range (0-255). Scale bars, 100 μm (A) and 50 μm (B and C).

## References and Notes

1. Coen, E., and Cosgrove, D.J. (2023). The mechanics of plant morphogenesis. Science 379, eade8055. 10.1126/science.ade8055.

2. Jones, M.A., Shen, J.J., Fu, Y., Li, H., Yang, Z., and Grierson, C.S. (2002). The Arabidopsis Rop2 GTPase is a positive regulator of both root hair initiation and tip growth. Plant Cell 14, 763–776. 10.1105/tpc.010359.

3. Molendijk, A.J., Bischoff, F., Rajendrakumar, C.S., Friml, J., Braun, M., Gilroy, S., and Palme, K. (2001). Arabidopsis thaliana Rop GTPases are localized to tips of root hairs and control polar growth. EMBO J 20, 2779–2788. 10.1093/emboj/20.11.2779.

4. Fu, Y., Gu, Y., Zheng, Z., Wasteneys, G., and Yang, Z. (2005). Arabidopsis interdigitating cell growth requires two antagonistic pathways with opposing action on cell morphogenesis. Cell 120, 687–700. 10.1016/j.cell.2004.12.026.

5. Settleman, J. (2005). Intercalating Arabidopsis leaf cells: a jigsaw puzzle of lobes, necks, ROPs, and RICs. Cell 120, 570–572. 10.1016/j.cell.2005.02.025.

6. Li, H., Lin, Y., Heath, R.M., Zhu, M.X., and Yang, Z. (1999). Control of pollen tube tip growth by a Rop GTPase-dependent pathway that leads to tip-localized calcium influx. Plant Cell 11, 1731–1742. 10.1105/tpc.11.9.1731.

7. Gu, Y., Fu, Y., Dowd, P., Li, S., Vernoud, V., Gilroy, S., and Yang, Z. (2005). A Rho family GTPase controls actin dynamics and tip growth via two counteracting downstream pathways in pollen tubes. J Cell Biol 169, 127–138. 10.1083/jcb.200409140.

8. Smokvarska, M., Jaillais, Y., and Martiniere, A. (2021). Function of membrane domains in rho-of-plant signaling. Plant Physiol 185, 663–681. 10.1093/plphys/kiaa082.

9. Ou, H., and Yi, P. (2022). ROP GTPase-dependent polarity establishment during tip growth in plants. New Phytol 236, 49–57. 10.1111/nph.18373.

10. Chang, F., Gu, Y., Ma, H., and Yang, Z. (2013). AtPRK2 promotes ROP1 activation via RopGEFs in the control of polarized pollen tube growth. Mol Plant 6, 1187–1201. 10.1093/mp/sss103.

11. Zhang, X., Jiang, H., Zhu, G., Chen, F., Yuan, W., Li, Y., Zhang, W., Wu, D., Liu, H., Wu, H.M., et al. (2025). RLCKs phosphorylate RopGEFs to control auxin-dependent Arabidopsis development. Nat Plants 11, 2130–2144. 10.1038/s41477-025-02111-9.

12. Bloch, D., Lavy, M., Efrat, Y., Efroni, I., Bracha-Drori, K., Abu-Abied, M., Sadot, E., and Yalovsky, S. (2005). Ectopic expression of an activated RAC in Arabidopsis disrupts membrane cycling. Mol Biol Cell 16, 1913–1927. 10.1091/mbc.e04-07-0562.

13. Cui, X., Wang, S., Huang, Y., Ding, X., Wang, Z., Zheng, L., Bi, Y., Ge, F., Zhu, L., Yuan, M., et al. (2022). Arabidopsis SYP121 acts as an ROP2 effector in the regulation of root hair tip growth. Mol Plant 15, 1008–1023. 10.1016/j.molp.2022.04.008.

14. Lavy, M., Bloch, D., Hazak, O., Gutman, I., Poraty, L., Sorek, N., Sternberg, H., and Yalovsky, S. (2007). A Novel ROP/RAC effector links cell polarity, root-meristem maintenance, and vesicle trafficking. Curr Biol 17, 947–952. 10.1016/j.cub.2007.04.038.

15. Lee, Y.J., Szumlanski, A., Nielsen, E., and Yang, Z. (2008). Rho-GTPase-dependent filamentous actin dynamics coordinate vesicle targeting and exocytosis during tip growth. J Cell Biol 181, 1155–1168. 10.1083/jcb.200801086.

16. Jones, M.A., Raymond, M.J., Yang, Z., and Smirnoff, N. (2007). NADPH oxidase-dependent reactive oxygen species formation required for root hair growth depends on ROP GTPase. J Exp Bot 58, 1261–1270. 10.1093/jxb/erl279.

17. Duan, Q., Kita, D., Li, C., Cheung, A.Y., and Wu, H.M. (2010). FERONIA receptor-like kinase regulates RHO GTPase signaling of root hair development. Proc Natl Acad Sci U S A 107, 17821–17826. 10.1073/pnas.1005366107.

18. Cosgrove, D.J. (2024). Plant Cell Wall Loosening by Expansins. Annu Rev Cell Dev Biol 40, 329–352. 10.1146/annurev-cellbio-111822-115334.

19. Cosgrove, D.J. (2023). Structure and growth of plant cell walls. Nat Rev Mol Cell Biol. 10.1038/s41580-023-00691-y.

20. Zhang, Y., Yu, J., Wang, X., Durachko, D.M., Zhang, S., and Cosgrove, D.J. (2021). Molecular insights into the complex mechanics of plant epidermal cell walls. Science 372, 706–711. 10.1126/science.abf2824.

21. Zhang, T., Tang, H., Vavylonis, D., and Cosgrove, D.J. (2019). Disentangling loosening from softening: insights into primary cell wall structure. Plant J 100, 1101–1117. 10.1111/tpj.14519.

22. Yu, J., Zhang, Y., and Cosgrove, D.J. (2024). The nonlinear mechanics of highly extensible plant epidermal cell walls. Proc Natl Acad Sci U S A 121, e2316396121. 10.1073/pnas.2316396121.

23. Wang, X., Wilson, L., and Cosgrove, D.J. (2020). Pectin methylesterase selectively softens the onion epidermal wall yet reduces acid-induced creep. J Exp Bot 71, 2629–2640. 10.1093/jxb/eraa059.

24. Denninger, P., Reichelt, A., Schmidt, V.A.F., Mehlhorn, D.G., Asseck, L.Y., Stanley, C.E., Keinath, N.F., Evers, J.F., Grefen, C., and Grossmann, G. (2019). Distinct RopGEFs Successively Drive Polarization and Outgrowth of Root Hairs. Curr Biol 29, 1854–1865 e1855. 10.1016/j.cub.2019.04.059.

25. Grierson, C., Nielsen, E., Ketelaarc, T., and Schiefelbein, J. (2014). Root hairs. Arabidopsis Book 12, e0172. 10.1199/tab.0172.

26. Bibikova, T.N., Jacob, T., Dahse, I., and Gilroy, S. (1998). Localized changes in apoplastic and cytoplasmic pH are associated with root hair development in Arabidopsis thaliana. Development 125, 2925–2934. 10.1242/dev.125.15.2925.

27. Steger, A., and Palmgren, M. (2022). Root hair growth from the pH point of view. Front Plant Sci 13, 949672. 10.3389/fpls.2022.949672.

28. Vissenberg, K., Fry, S.C., and Verbelen, J.P. (2001). Root hair initiation is coupled to a highly localized increase of xyloglucan endotransglycosylase action in arabidopsis roots. Plant Physiology 127, 1125–1135. DOI 10.1104/pp.010295.

29. Schoenaers, S., Lee, H.K., Gonneau, M., Faucher, E., Levasseur, T., Akary, E., Claeijs, N., Moussu, S., Broyart, C., Balcerowicz, D., et al. (2024). Rapid alkalinization factor 22 has a structural and signalling role in root hair cell wall assembly. Nat Plants 10, 494–511. 10.1038/s41477-024-01637-8.

30. Zhou, K., Hepler, N.K., Jia, M., and Cosgrove, D.J. (2026). Functional divergence of plant expansins as revealed by genetic complementation of root-hair tip growth in Arabidopsis thaliana. Proc Natl Acad Sci U S A 123, e2623483123. 10.1073/pnas.2623483123.

31. McQueen-Mason, S., and Cosgrove, D.J. (1994). Disruption of hydrogen bonding between plant cell wall polymers by proteins that induce wall extension. Proc Natl Acad Sci U S A 91, 6574–6578. 10.1073/pnas.91.14.6574.

32. Grebe, M., Friml, J., Swarup, R., Ljung, K., Sandberg, G., Terlou, M., Palme, K., Bennett, M.J., and Scheres, B. (2002). Cell polarity signaling in Arabidopsis involves a BFA-sensitive auxin influx pathway. Curr Biol 12, 329–334. 10.1016/s0960-9822(02)00654-1.

33. Sieberer, T., Seifert, G.J., Hauser, M.T., Grisafi, P., Fink, G.R., and Luschnig, C. (2000). Post-transcriptional control of the Arabidopsis auxin efflux carrier EIR1 requires AXR1. Curr Biol 10, 1595–1598. 10.1016/s0960-9822(00)00861-7.

34. Luschnig, C., and Friml, J. (2024). Over 25 years of decrypting PIN-mediated plant development. Nat Commun 15, 9904. 10.1038/s41467-024-54240-y.

35. Zhuang, X., Li, R., and Jiang, L. (2024). A century journey of organelles research in the plant endomembrane system. Plant Cell 36, 1312–1333. 10.1093/plcell/koae004.

36. Sztul, E., Chen, P.W., Casanova, J.E., Cherfils, J., Dacks, J.B., Lambright, D.G., Lee, F.S., Randazzo, P.A., Santy, L.C., Schurmann, A., et al. (2019). ARF GTPases and their GEFs and GAPs: concepts and challenges. Mol Biol Cell 30, 1249–1271. 10.1091/mbc.E18-12-0820.

37. Mohr, I., Sanyal, S.K., Ivanov, R., and Bauer, P. (2026). The plant ARF GTPase family: current perspectives in membrane trafficking, cellular signaling, and new paradigms. Plant Cell Physiol 67, 1677–1692. 10.1093/pcp/pcag079.

38. Geldner, N., Anders, N., Wolters, H., Keicher, J., Kornberger, W., Muller, P., Delbarre, A., Ueda, T., Nakano, A., and Jurgens, G. (2003). The Arabidopsis GNOM ARF-GEF mediates endosomal recycling, auxin transport, and auxin-dependent plant growth. Cell 112, 219–230. 10.1016/s0092-8674(03)00003-5.

39. Gendre, D., McFarlane, H.E., Johnson, E., Mouille, G., Sjodin, A., Oh, J., Levesque-Tremblay, G., Watanabe, Y., Samuels, L., and Bhalerao, R.P. (2013). Trans-Golgi network localized ECHIDNA/Ypt interacting protein complex is required for the secretion of cell wall polysaccharides in Arabidopsis. Plant Cell 25, 2633–2646. 10.1105/tpc.113.112482.

40. Renna, L., Stefano, G., Slabaugh, E., Wormsbaecher, C., Sulpizio, A., Zienkiewicz, K., and Brandizzi, F. (2018). TGNap1 is required for microtubule-dependent homeostasis of a subpopulation of the plant trans-Golgi network. Nat Commun 9, 5313. 10.1038/s41467-018-07662-4.

41. Gendre, D., Baral, A., Dang, X., Esnay, N., Boutte, Y., Stanislas, T., Vain, T., Claverol, S., Gustavsson, A., Lin, D., et al. (2019). Rho-of-plant activated root hair formation requires Arabidopsis YIP4a/b gene function. Development 146. 10.1242/dev.168559.

42. Fu, Y., Xu, T., Zhu, L., Wen, M., and Yang, Z. (2009). A ROP GTPase signaling pathway controls cortical microtubule ordering and cell expansion in Arabidopsis. Curr Biol 19, 1827–1832. 10.1016/j.cub.2009.08.052.

43. Xiang, X., Zhang, S., Li, E., Shi, X.L., Zhi, J.Y., Liang, X., Yin, G.M., Qin, Z., Li, S., and Zhang, Y. (2023). RHO OF PLANT proteins are essential for pollen germination in Arabidopsis. Plant Physiol 193, 140–155. 10.1093/plphys/kiad196.

44. Fu, Y., Li, H., and Yang, Z. (2002). The ROP2 GTPase controls the formation of cortical fine F-actin and the early phase of directional cell expansion during Arabidopsis organogenesis. Plant Cell 14, 777–794. 10.1105/tpc.001537.

45. Mulvey, H., and Dolan, L. (2025). The evolution of RHO of plant (ROP) proteins and their morphogenetic functions. New Phytol 248, 542–548. 10.1111/nph.70455.

46. Feiguelman, G., Fu, Y., and Yalovsky, S. (2018). ROP GTPases Structure-Function and Signaling Pathways. Plant Physiol 176, 57–79. 10.1104/pp.17.01415.

47. Gendre, D., Oh, J., Boutte, Y., Best, J.G., Samuels, L., Nilsson, R., Uemura, T., Marchant, A., Bennett, M.J., Grebe, M., and Bhalerao, R.P. (2011). Conserved Arabidopsis ECHIDNA protein mediates trans-Golgi-network trafficking and cell elongation. Proc Natl Acad Sci U S A 108, 8048–8053. 10.1073/pnas.1018371108.

48. Hachez, C., Laloux, T., Reinhardt, H., Cavez, D., Degand, H., Grefen, C., De Rycke, R., Inze, D., Blatt, M.R., Russinova, E., and Chaumont, F. (2014). Arabidopsis SNAREs SYP61 and SYP121 coordinate the trafficking of plasma membrane aquaporin PIP2;7 to modulate the cell membrane water permeability. Plant Cell 26, 3132–3147. 10.1105/tpc.114.127159.

49. Oda, Y., and Fukuda, H. (2012). Initiation of cell wall pattern by a Rho- and microtubule-driven symmetry breaking. Science 337, 1333–1336. 10.1126/science.1222597.

50. Zheng, Z.L., Nafisi, M., Tam, A., Li, H., Crowell, D.N., Chary, S.N., Schroeder, J.I., Shen, J., and Yang, Z. (2002). Plasma membrane-associated ROP10 small GTPase is a specific negative regulator of abscisic acid responses in Arabidopsis. Plant Cell 14, 2787–2797. 10.1105/tpc.005611.

51. Lin, W., Tang, W., Pan, X., Huang, A., Gao, X., Anderson, C.T., and Yang, Z. (2022). Arabidopsis pavement cell morphogenesis requires FERONIA binding to pectin for activation of ROP GTPase signaling. Curr Biol 32, 497–507 e494. 10.1016/j.cub.2021.11.030.

52. Xu, T., Dai, N., Chen, J., Nagawa, S., Cao, M., Li, H., Zhou, Z., Chen, X., De Rycke, R., Rakusova, H., et al. (2014). Cell surface ABP1-TMK auxin-sensing complex activates ROP GTPase signaling. Science 343, 1025–1028. 10.1126/science.1245125.

53. Li, E., Zhang, Y.L., Shi, X., Li, H., Yuan, X., Li, S., and Zhang, Y. (2021). A positive feedback circuit for ROP-mediated polar growth. Mol Plant 14, 395–410. 10.1016/j.molp.2020.11.017.

54. Xu, J., and Scheres, B. (2005). Dissection of Arabidopsis ADP-RIBOSYLATION FACTOR 1 function in epidermal cell polarity. Plant Cell 17, 525–536. 10.1105/tpc.104.028449.

55. Grefen, C., Donald, N., Hashimoto, K., Kudla, J., Schumacher, K., and Blatt, M.R. (2010). A ubiquitin-10 promoter-based vector set for fluorescent protein tagging facilitates temporal stability and native protein distribution in transient and stable expression studies. Plant J 64, 355–365. 10.1111/j.1365-313X.2010.04322.x.

56. Cutler, S.R., Ehrhardt, D.W., Griffitts, J.S., and Somerville, C.R. (2000). Random GFP::cDNA fusions enable visualization of subcellular structures in cells of Arabidopsis at a high frequency. Proc Natl Acad Sci U S A 97, 3718–3723. 10.1073/pnas.97.7.3718.

